# Ribosome collision sensor Hel2 recognizes mistargeting secretory ribosome-nascent chain complexes

**DOI:** 10.1101/2020.12.28.424499

**Authors:** Yoshitaka Matsuo, Toshifumi Inada

## Abstract

Ribosome collision due to translational stalling is recognized as a problematic event in translation by E3 ubiquitin ligase Hel2, leading to non-canonical subunit dissociation followed by targeting of the faulty nascent peptides for degradation. Although Hel2-mediated quality control greatly contributes to maintaining cellular protein homeostasis, its physiological role in dealing with endogenous substrates remains unclear. This study utilized genome-wide analysis, based on selective ribosome profiling, to survey the endogenous substrates for Hel2. This survey revealed that Hel2 preferentially binds to the pre-engaged secretory ribosome-nascent-chain complexes (RNCs), which translate upstream of targeting signals. Notably, Hel2 recruitment into secretory RNCs was elevated under signal recognition particle (SRP)-deficient conditions. Moreover, the mitochondrial defects caused by insufficient SRP were enhanced by *hel2* deletion, along with the mistargeting of secretory proteins into mitochondria. Collectively, these findings provide novel insights into risk management in the secretory pathway that maintains cellular protein homeostasis.

## Introduction

Protein biosynthesis on ribosomes can fail for numerous reasons, including genetic mutations, mRNA processing errors, lack of availability of charged tRNAs, and the properties of nascent protein chains (Brandman and Hegde, 2016; Collart and Weiss, 2019; Joazeiro, 2019). Because these products may function anomalously, such as by forming toxic aggregates (Chiti and Dobson, 2017; Choe et al., 2016; Yonashiro et al., 2016), surveillance systems within cells monitor every step of translation and dispose of such products to prevent their accumulation (Brandman and Hegde, 2016; Collart and Weiss, 2019; Joazeiro, 2019).

Because elongation of translation is tightly associated with biological processes, such as co-translational protein folding (Doring et al., 2017; Kramer et al., 2009; Oh et al., 2011; Pechmann and Frydman, 2013; Stein et al., 2019), protein targeting (Chartron et al., 2016; Pechmann et al., 2014; Schibich et al., 2016), protein assembly (Panasenko et al., 2019; Shiber et al., 2018), and mRNA decay (Buschauer et al., 2020; Presnyak et al., 2015), the speed of translation of each coding region of mRNA is individually regulated to produce fully functional proteins. Thus, normal translation speeds range widely in the cellular translation pool (Collart and Weiss, 2019; Stein and Frydman, 2019). Accordingly, the selectivity for aberrant and harmful ribosome stalling is determined by ribosome queuing rather than simply by translation slowdown to avoid targeting pausing of programmed ribosomes (Ikeuchi et al., 2019; Juszkiewicz et al., 2018; Simms et al., 2017). This mechanism can identify many types of aberrant ribosome stalling in diverse translational pools.

For quality control, the ubiquitin ligase Hel2 recognizes ribosome collision as a problematic event in translation and is responsible for the initiation of two different pathways, ribosome-associated quality control (RQC) and No-Go mRNA decay (NGD). In the RQC pathway, Hel2 ubiquitinates ribosomal protein uS10 to initiate non-canonical subunit dissociation, followed by targeting of the faulty nascent peptides for degradation (Bengtson and Joazeiro, 2010; Brandman et al., 2012; Matsuo et al., 2017; Shen et al., 2015). A similar mechanism is conserved in mammalian cells (Juszkiewicz and Hegde, 2016; Sundaramoorthy et al., 2017). In yeast, Hel2 also induces NGD via a Cue2-mediated endonucleolytic cleavage event (D’Orazio et al., 2019; Ikeuchi et al., 2019; Simms et al., 2017). Furthermore, the human Hel2 homolog ZNF598 was shown to inhibit translation initiation at the ribosome collision site by the GIGYF2-4EHP system in mammalian cells (Hickey et al., 2020). Although 4EHP is not conserved in yeast, the yeast GIGYF2 homologs Smy2 and Syh1 mediate mRNA decay at ribosome collision sites in yeast (Hickey et al., 2020), indicating that Hel2 acts as a master regulator to determine the fate of the colliding ribosomes.

The colliding ribosomes recognized by Hel2/ZNF598 are not merely juxtaposed on the same mRNA; rather, this provides the unique structural architecture at the collision interface (Ikeuchi et al., 2019; Juszkiewicz et al., 2018; Juszkiewicz et al., 2020; Matsuo et al., 2020), which can serve as a scaffold for Hel2/ZNF598-mediated reactions (Winz et al., 2019). However, the Hel2 position on the colliding ribosomes remains to be elucidated. Although ribosome profiling techniques including disome profiling can detect ribosomal collisions in endogenous genes, the detectable collision events are mainly linked to other biological processes, such as protein folding and protein targeting, rather than to quality control (Arpat et al., 2020 Guydosh and Green, 2014; Han et al., 2019; Meydan and Guydosh, 2020; Zhao et al., 2020). Moreover, the unique structural architecture, which forms at the interface of aberrantly colliding ribosomes, is not observable in natural disome structures prepared from rapidly proliferating cells (Zhao et al., 2020). Thus, although Hel2-triggered quality control could greatly contribute to maintaining cellular protein homeostasis, its physiological role in dealing with endogenous substrates remains unclear.

This study presents a genome-wide analysis, based on selective ribosome profiling, to identify the endogenous substrates of Hel2. This straightforward approach showed that Hel2 frequently recognized aberrant ribosome collisions on secretory mRNAs prior to endoplasmic reticulum (ER) engagement. In the initial selection of secretory ribosome-nascent-chain complexes (RNCs) by signal recognition particles (SRP), Hel2 recognizes the harmful substrates lacking SRP recognition. This triage system greatly contributes to preventing the mistargeting of secretory proteins into mitochondria. These findings provide novel insights into how Hel2 contributes to risk management in the secretory pathway that maintains cellular protein homeostasis.

## Results

### Selective ribosome profiling for Hel2-associated colliding ribosomes

To address the physiological role of the Hel2-mediated quality control system, we sought to identify the endogenous targets of Hel2 using a genome-wide approach based on selective ribosome profiling (Becker et al., 2013; Galmozzi et al., 2019; Ingolia et al., 2009). Previous cryo-EM analysis showed that the vast majority of Hel2-associated ribosomes are occupied with mRNAs (Matsuo et al., 2017). Endogenous Hel2-associated ribosomes were affinity purified from a yeast strain in which chromosomal Hel2 was tagged with Flag-TEV-ProteinA tag (FTP), followed by sequencing of the ribosome-protected mRNA fragments (Hel2-IPed ribo-seq) **(Figure 1A)**.

**Figure 1.**
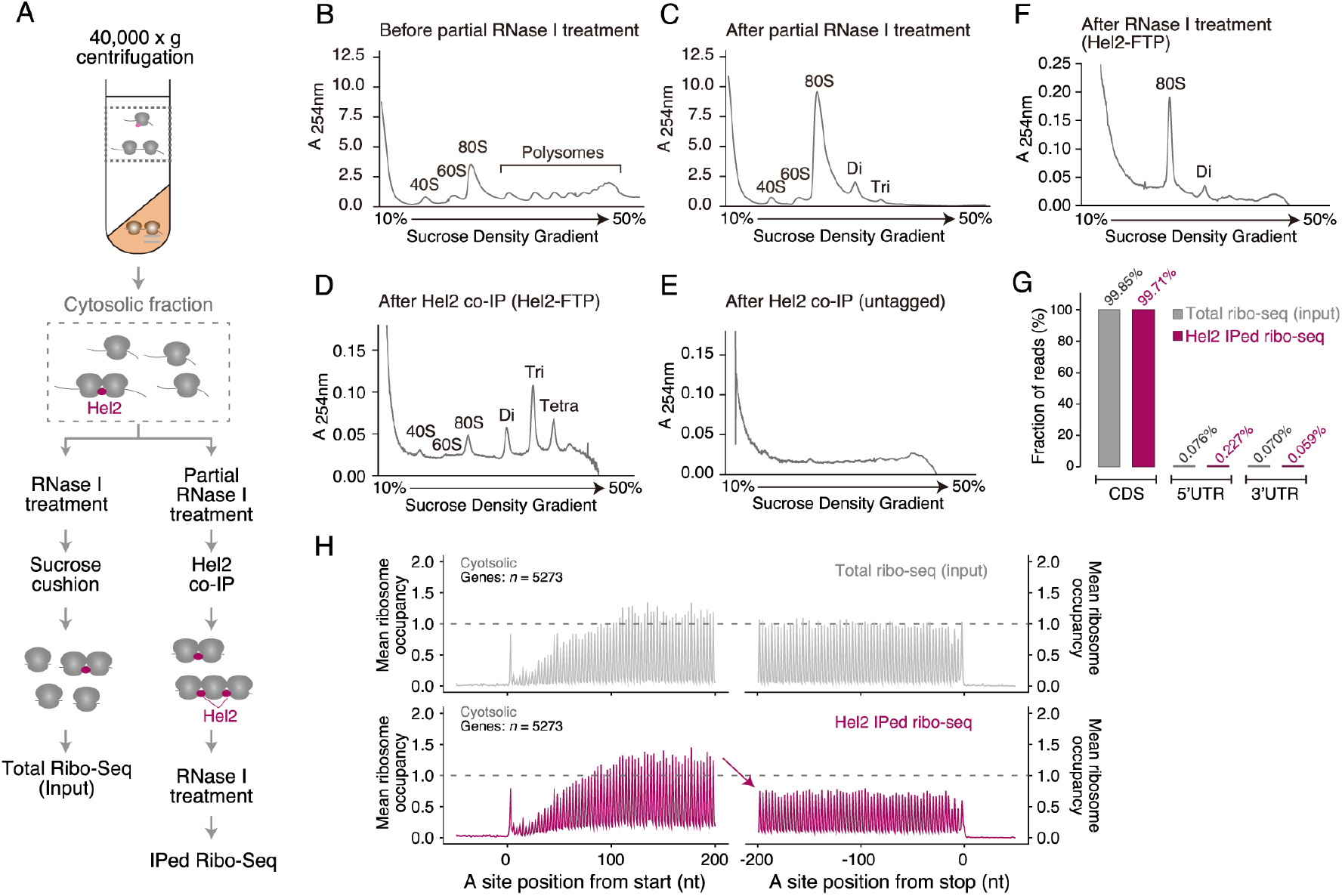
Selective ribosome profiling for Hel2-associated ribosomes. **(A)** Experimental setup of selective ribosome profiling. Hel2-associated ribosomes and total ribosomes were prepared by immunoprecipitation and sucrose cushion centrifugation, respectively. **(B–F)** Sucrose density gradient sedimentation analysis. The disrupted cell powder was resuspended in lysis buffer and centrifuged at 40,000 x g for 30 minutes at 4 °C **(B)**. The supernatant was partially digested with RNase I for 1 hour at 4 °C **(C)**, and Hel2-associated ribosomes were purified by immunoprecipitation **(D)**. The supernatant prepared from untagged-yeast strain was used as a negative control for immunoprecipitation **(E)**. The purified Hel2-associated ribosomes were digested with RNase I for 45 minutes at 23°C to generate ribosome-protected mRNA fragments **(F)**. The fraction in each step of Hel2-associated ribosome purification was analyzed by 10–50% (w/v) sucrose density gradient centrifugation. **(G)** Fraction of the reads mapping to each reading frame in the coding sequence (CDS), the 5’ untranslated region (UTR), and the 3’ UTR for total and Hel2-IPed ribo-seq. **(H)** Metagene plot of ribosome occupancy. The ribosome occupancy score was calculated as the ratio of reads at a given nt position to average reads per codon on the individual ORF (**Figure S2C**). Average ribosome occupancy plotted around the start and stop codons at the A site for total ribo-seq and Hel2-IPed ribo-seq. The 28 nt reads and the transcripts containing more than 0.5 reads per codon were analyzed. The offset was 15.

During purification of Hel2-associated ribosomes, two translation elongation inhibitors, cycloheximide (CHX) and tigecycline (TIG), were introduced into the lysis buffer to prevent ribosome movement. To ensure the selectivity of Hel2-associated ribosomes, mRNAs were partially digested with RNase I at 4 °C before purification, separating the individual ribosome engaged on the same mRNA (**Figure 1B-C**). Consistent with previous findings (Ikeuchi et al., 2019; Juszkiewicz et al., 2018; Juszkiewicz et al., 2020; Matsuo et al., 2020), the purified Hel2-associated ribosomes formed multimeric-ribosome complexes, including di-, tri-, and tetra-somes, even after partial RNase I treatment (**Figure 1D**), suggesting that Hel2 indeed binds to endogenous colliding ribosomes. By contrast, no ribosomes were purified using untagged-yeast strain (**Figure 1E**). Although the tightly packaged colliding ribosomes on the reporter mRNA containing a very strong artificial arrest sequence was nuclease resistant (Juszkiewicz et al., 2018), the purified endogenous Hel2-associated colliding di-, tri-, and tetra-somes were digested to monosomes following a second treatment with excess RNase I at 23 °C, which also generated ribosome-protected mRNA fragments (**Figure 1F**). Thus, we further examined the RNase resistance level of representative Hel2-substrates on the SDD1 mRNA against second RNase treatment using a previously established in vitro reconstitution system (Matsuo et al., 2020). As expected, this clearly showed that Hel2-targeting colliding ribosomes including di-, tri-, and tetra-somes on the SDD1 mRNAs were completely converted to monosomes after the same treatment as the second RNase I treatment (Figure S1). Although it is not a precise determination for the number of ribosomes involved in the queue and which ribosome interacts with Hel2, we concluded endogenous Hel2-associated colliding ribosomes could be recovered as monosomes in this system. Therefore, we here focused on the monosome footprints after the second RNase I treatment for Hel2-IPed ribo-seq.

In parallel, a total ribosome fraction from the same lysate was prepared by sucrose cushioning (**Figure 1A**), generating a total ribosome profiling library (total ribo-seq: input). The ribosome occupancy score and Hel2-enrichment score for individual coding sequences were calculated using two datasets: Hel2-IPed ribo-seq and total ribo-seq **(Figure S2A and S2B)**. The ribosome occupancy score, a measure of ribosome density on individual coding sequences of each dataset, was calculated as the ratio of reads at a given codon to average reads per codon on an individual open reading frame (ORF) (**Figure S2C**). The Hel2-enrichment score at codon resolution was calculated by dividing the reads per million (RPM) of Hel2-IPed ribo-seq by RPM of total ribo-seq (**Figure S2C**).

The footprints derived from Hel2-IPed ribo-seq were almost completely mapped to the coding sequence to a similar extent as total ribo-seq (input) (**Figure 1G**). Metagene analysis, defined as the mean ribosome occupancy score of overall transcripts (n = 5273) around start and stop codons at nucleotide resolution, showed that Hel2-associated ribosome occupancy was significantly increased at the 5’ region of transcripts, but was decreased at the 3’ region (**Figure 1H)**, indicating that Hel2 was apparently associated with ribosomes during the early phase of translation elongation.

### Hel2 preferentially binds to secretory RNCs

To further investigate the functional properties of Hel2 targets, Hel2-enrichment scores of overall transcripts (n = 5276) were calculated using R package “DEseq2” (Love et al., 2014). A subset of Hel2-enriched mRNAs with Hel2-enrichment (log2) >2 and P_adj_ <0.01 was chosen **(Figure 2A)**. The Hel2-enriched mRNAs (n = 602) were subjected to gene ontology (GO) enrichment analysis (http://geneontology.org). Analysis of the cellular component showed that Hel2-associated ribosomes were highly enriched in transcripts encoding membrane proteins **(Figure 2B)**, but were significantly depleted of transcripts encoding cytosolic protein **(Figure 2B)**. Many transcripts encoding membrane protein are co-translationally associated with ER, resulting in the translocation of nascent chains into the membrane (Guna and Hegde, 2018; Jan et al., 2014; Zhang and Shan, 2014). Therefore, to assess the specificity of Hel2 targets for ER-targeted transcripts, Hel2 enrichment of overall transcripts were compared among several cellular compartments. Notably, significant Hel2-enrichment was observed in ER-targeted mRNAs containing a signal sequence (SS) or transmembrane domain (TMD) coding sequences **(Figure 2C)**; similar enrichment was not observed in mRNAs encoding nuclear, cytoplasmic, mitochondrial, and tail-anchored proteins associated post-translationally with membranes (Guna and Hegde, 2018; Kutay et al., 1995) **(Figure 2C)**. Further investigation of Hel2-enrichment in previously annotated secretory mRNAs with expected SRP dependence (Ast et al., 2013; Chartron et al., 2016; Costa et al., 2018) showed that Hel2 apparently binds to all types of mRNAs encoding secretory proteins, including SRP-dependent, SRP-independent, and internal proteins (**Figure 2D**). Collectively, these findings indicate that the characteristic feature of Hel2 targets was co-translational engagement with the ER rather than membrane proteins.

**Figure 2.**
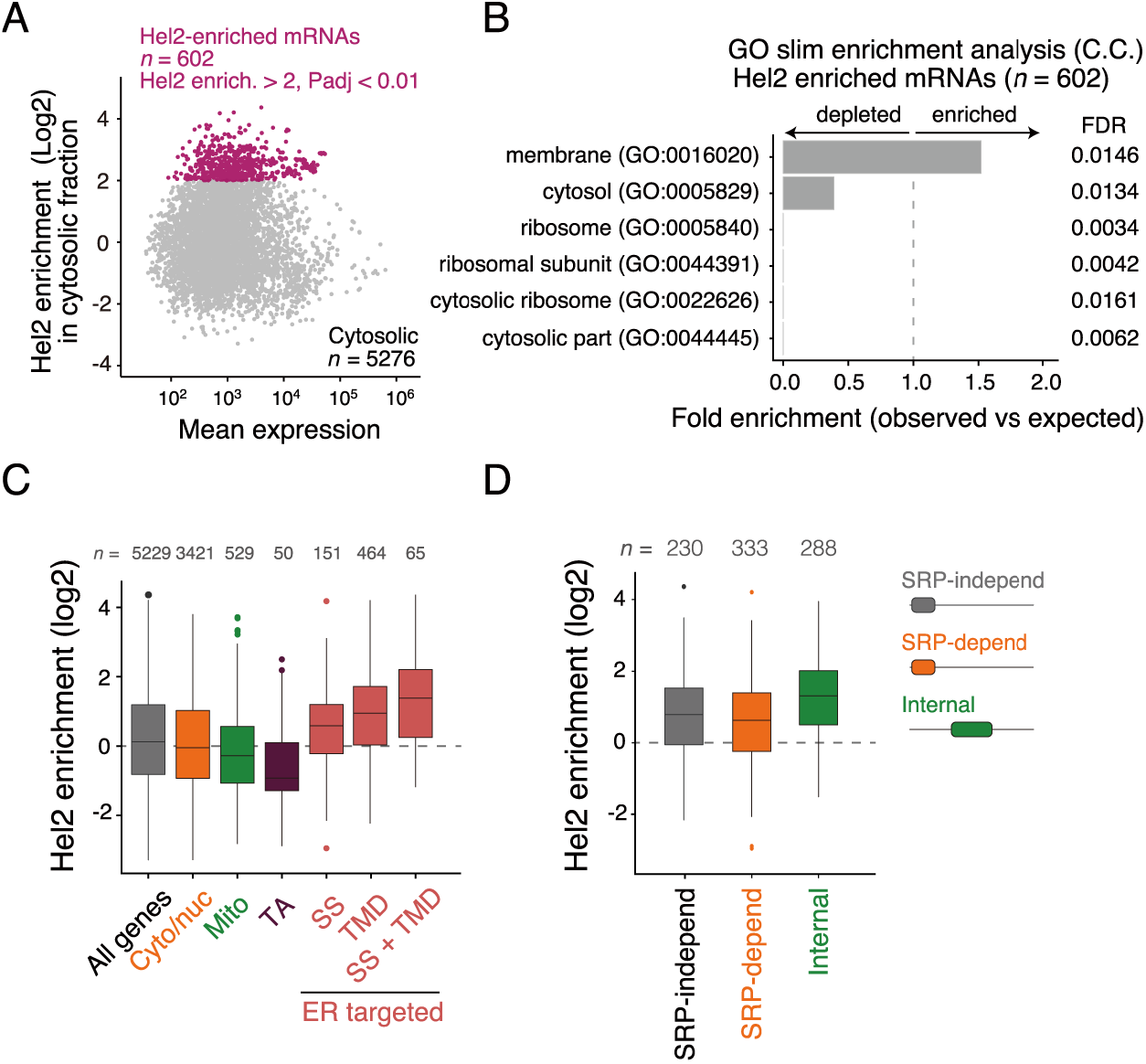
Hel2 preferentially binds to secretory RNCs. **(A)** Scatter plot of Hel2 enrichment, expressed as fold change between Hel2-IPed ribo-seq and total ribo-seq, versus expression level in total ribo-seq. Hel2 enrichment scores at individual transcripts were calculated by R package DEseq2. Analyses have restricted the transcripts mapped with more than 25 reads. Highlighted Hel2-enriched mRNAs were defined as those with Hel2 enrichment scores (log2) > 2 and P_adj_ < 0.01. P-values and P_adj_ were calculated by the Wald test and the Benjamini-Hochberg method, respectively. All Hel2-enrichment scores and P_adj_ for individual transcripts are provided in Supplemental Table 1. **(B)** Gene ontology enrichment analysis. Hel2 enriched transcripts indicated in (A) were analyzed by GO enrichment analysis in cellular component at (http://geneontology.org). The GO terms defined by those with < 0.05 FDR (False Discovery Rate) were displayed in the panel. **(C)** Box plot of Hel2 enrichment scores of overall transcripts among cellular compartments. Classification of individual transcripts to cellular compartments was as described (Chartron et al., 2016). **(D)** Box plot of Hel2 enrichment scores of the transcripts encoding secretory proteins. Classification of individual transcripts as being SRP dependence has been described (Costa et al., 2018). The first TMD of internal proteins was after the initial 60 amino acids.

### Hel2 binds to secretory RNCs that are not yet engaged with ER

Because most secretory proteins are co-translationally translocated into and across the ER membrane, only a small fraction of their transcripts are translated on the cytosolic ribosome before engagement with the ER (Guna and Hegde, 2018; Zhang and Shan, 2014). The initial procedure for the purification of Hel2-associated ribosomes did not allow solubilization of the membrane fraction, suggesting that Hel2 could potentially target pre- or mis-engaged transcripts associated with the ER. To confirm this possibility, cells were lysed with 1% Triton X-100 to solubilize membrane-associated mRNAs **(Figure 3A)**. Monitoring ribosome status during purification steps by sucrose gradient sedimentation showed that 1% Triton X-100 had no effect on the purification of Hel2-associated ribosomes (**Figure S3A-E**). In addition, comparisons of total ribo-seq footprints in the cytosolic and membrane-solubilized fraction showed that ER-targeted mRNAs were indeed solubilized by 1% Triton X-100 treatment **(Figure S4A-B)**. Thus, the membrane-solubilized fraction prepared by treatment with 1% Triton X-100 was subjected to selective ribosome profiling (**Figure S3F-G**). In the membrane-solubilized fraction, the footprints derived from Hel2-associated ribosomes were again completely mapped to the coding sequence (**Figure S3H-I**), and the Hel2-associated ribosome density was higher at the 5’ than at the 3’ region (**Figure S3I**). However, the Hel2-enrichment of ER-targeted mRNAs, which is monitored in the cytosolic fraction, totally disappeared **(Figure S4C-D)**, indicating that the solubilized membrane-associated RNCs were not targeted by Hel2. These findings indicated that Hel2 frequently targeted pre-engaged, but not post-engaged, secretory RNCs.

**Figure 3.**
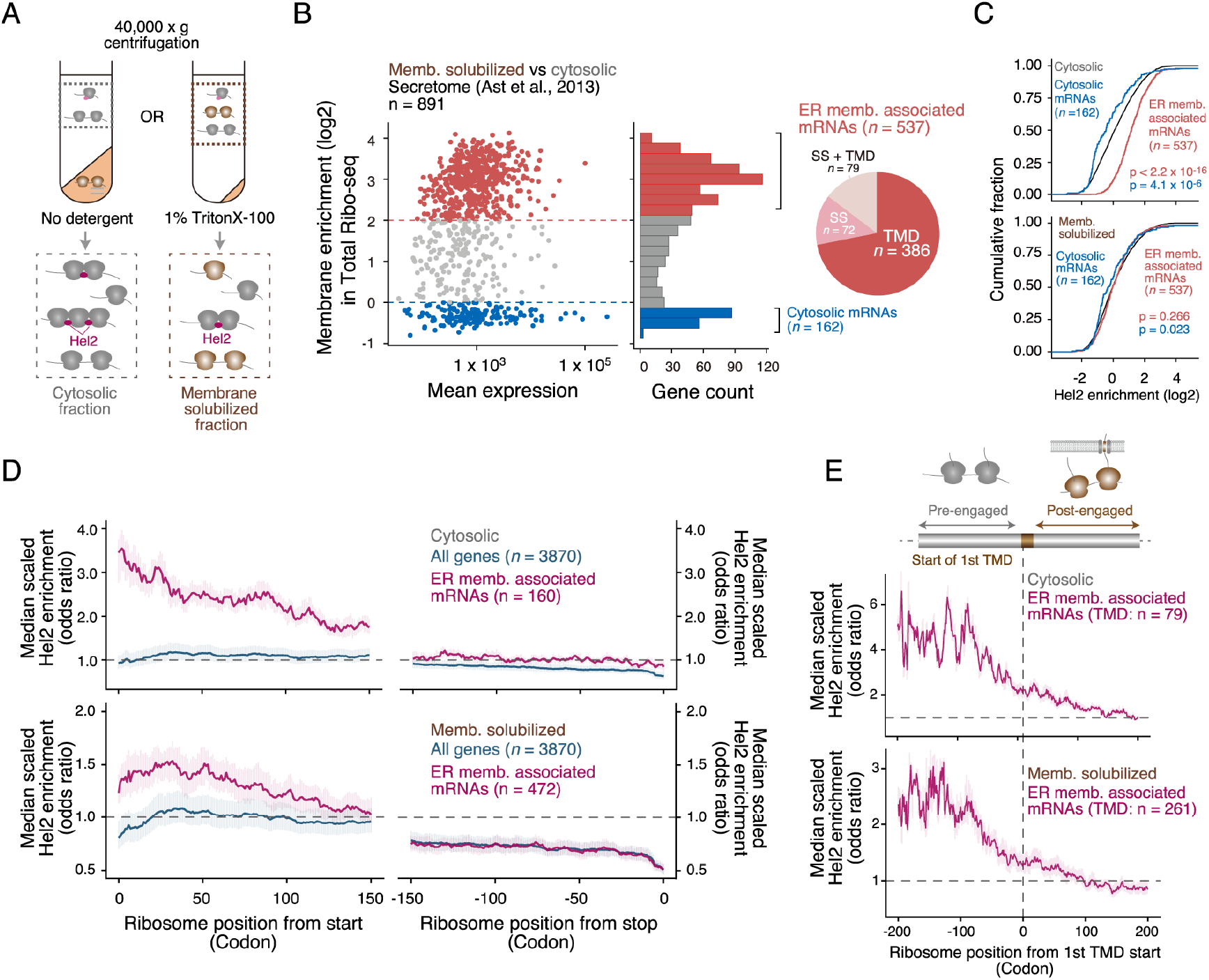
Hel2 monitors the translational status of pre-engaged secretory RNCs. **(A)** Experimental set up of membrane solubilization. Membrane-associated RNCs were obtained by the membrane solubilization with 1% Triton X-100. **(B)** Left panel: Scatter plot of membrane enrichment in total ribo-seq, expressed as fold change between membrane-solubilized and cytosolic fractions versus mean expression level in total ribo-seq. Membrane enrichment scores of individual ER targeted mRNAs (Ast et al., 2013) were calculated by R package DEseq2. Analyses have restricted the transcripts mapped with more than 25 reads. Middle panel: Histogram of membrane enrichments. The bin width was 0.25. ER membrane-associated mRNAs (Red) were defined as those with (log2) > 2 membrane enrichment and P_adj_ < 0.01. Cytosolic mRNAs (Blue) were defined by those with (log2) < 0 membrane enrichment. P-values and P_adj_ were calculated using the Wald test and the Benjamini-Hochberg method, respectively. Right panel: Classification of ER membrane-associated mRNAs. **(C)** Cumulative distribution of Hel2 enrichment obtained with all transcripts (gray), ER membrane-associated mRNAs (red), and cytosolic mRNAs (Blue). P-values were calculated by Student’s t-tests. **(D)** Metagene plot of Hel2 enrichment score. Hel2 enrichment scores were calculated by dividing the RPM of Hel2-IPed ribo-seq by the RPM of total ribo-seq (**Figure S2C**). Median-scaled Hel2 enrichment scores of all genes or ER membrane-associated mRNAs, defined in (B), were plotted around the start and stop codons for the cytosolic and membrane-solubilized fractions. Transcripts containing more than 0.5 reads per codon were further analyzed. **(E)** Metagene plot of Hel2 enrichment score. Median-scaled Hel2 enrichment scores of ER membrane-associated mRNAs encoding internal proteins were plotted around the first TMD coding region (Chartron et al., 2016) for the cytosolic and membrane-solubilized fractions. Transcripts containing more than 0.5 reads per codon were further analyzed.

### Hel2 monitors the initial selection step for the secretory pathway

We employed the classification of the ER-targeted mRNAs, which are derived from the secretory proteins previously predicted to translocate into the ER (Ast et al., 2013). However, some of these mRNAs were translated in the cytosol rather than on the ER membrane **(Figure 3B)**. These mRNAs were therefore redefined as ER membrane-associated mRNAs (log2 membrane enrichment > 2: n = 537) and cytosolic mRNAs (log2 membrane enrichment < 0: n = 162) based on the degree of membrane enrichment (**Figure 3B**). This classification allowed us to focus specifically on ER membrane-associated mRNAs, which were clearly targeted by Hel2 **(Figure 3C)**.

Co-translational targeting to the ER mainly depends on the SRP (Ast et al., 2013; Guna and Hegde, 2018; Zhang and Shan, 2014). In the classical model, SRP is recruited to the secretory RNCs by the targeting signal of the nascent peptide (Guna and Hegde, 2018; Zhang and Shan, 2014). However, a recent study showed that SRP can be recruited to secretory RNCs before the targeting signal is accessible and can facilitate translocation only after the emergence of a targeting signal from the ribosome tunnel (Chartron et al., 2016). This SRP-preloading model implies that the selection of RNCs for protein targeting in the secretory pathway is initiated even before the targeting signal is translated. Because Hel2 preferentially binds to secretory RNCs before ER engagement, we hypothesized that Hel2 might monitor the initial selection step in the secretory pathway.

To test this hypothesis, we investigated the distribution of Hel2-associated ribosomes within the ORF of ER membrane-associated mRNAs by meta-analysis, in which median-scaled Hel2-enrichment scores were plotted around the start and stop codons. In the cytosolic fraction, Hel2-associated ribosomes showed marked accumulation at the 5’ region, but not at the 3’ region, of the ORF (**Figure 3D**). This tendency was observed only in ER membrane-associated mRNAs, not in all genes or in other subsets, including cytonuclear and mitochondrial genes (**Figure 3D & S4E**). The accumulation of Hel2-associated ribosomes at the 5’ region was also observed in the membrane-solubilized fraction, but to a lesser extent than in the cytosolic fraction (**Figure 3D**), whereas the accumulation of Hel2-associated ribosomes at the 3’ region of the membrane-solubilized fraction was significantly depleted (**Figure 3D**). To further investigate the Hel2 distribution of RNCs before and after engagement with the ER, we focused on the TMD coding mRNAs and analyzed Hel2-enrichment around the first TMD coding region. Hel2-associated ribosomes were again highly enriched at the area upstream of the first TMD coding region in both cytosolic and membrane-solubilized fractions (**Figure 3E)**. These results indicated that Hel2 mainly binds to pre-engaged secretory RNCs translating upstream of the targeting signal, supporting our hypothesis that Hel2 monitors the initial selection step in the secretory pathway.

The translating ribosomes on the secretory mRNAs tend to slow down downstream of the signal targeting membrane engagement (Pechmann et al., 2014), suggesting that this slow down may be sufficient to generate colliding ribosomes for Hel2 recruitment. To carefully monitor ribosome density around the targeting signal, we focused on the top-ranked Hel2-enriched internal proteins (**Figure 4A**; n = 45). The TMD coding region, which acts as the targeting signal in these internal proteins, is located more than 60 codons downstream of the start codon. Then we calculated median-scaled ribosome occupancy in the total ribo-seq dataset (input). In the cytosolic fraction, the ribosome density (input) began to decrease after the first TMD start site, with ribosome occupancy being below average after emergence of the first TMD from the ribosome tunnel (**Figure 4B**: left upper panel), indicating that these RNCs engage with the ER membrane after exposure of the first TMD. By contrast, the translating ribosomes in the membrane-solubilized fraction were evenly distributed across the entire region up- and downstream of the first TMD coding region (**Figure 4B**: right upper panel), which contained the membrane-associated RNCs. Calculation of the median-scaled ribosome occupancy in the Hel2-IPed ribo-seq dataset revealed that the patterns of Hel2-associated ribosome density were similar to the pattern of cytosolic fraction, with ribosome density being highly concentrated upstream of the first TMD coding region (**Figure 4B-C**). In addition, Hel2 recruitment was observed upstream of the first TMD coding region, not as a peak just downstream of that region (**Figure 4B**).

**Figure 4.**
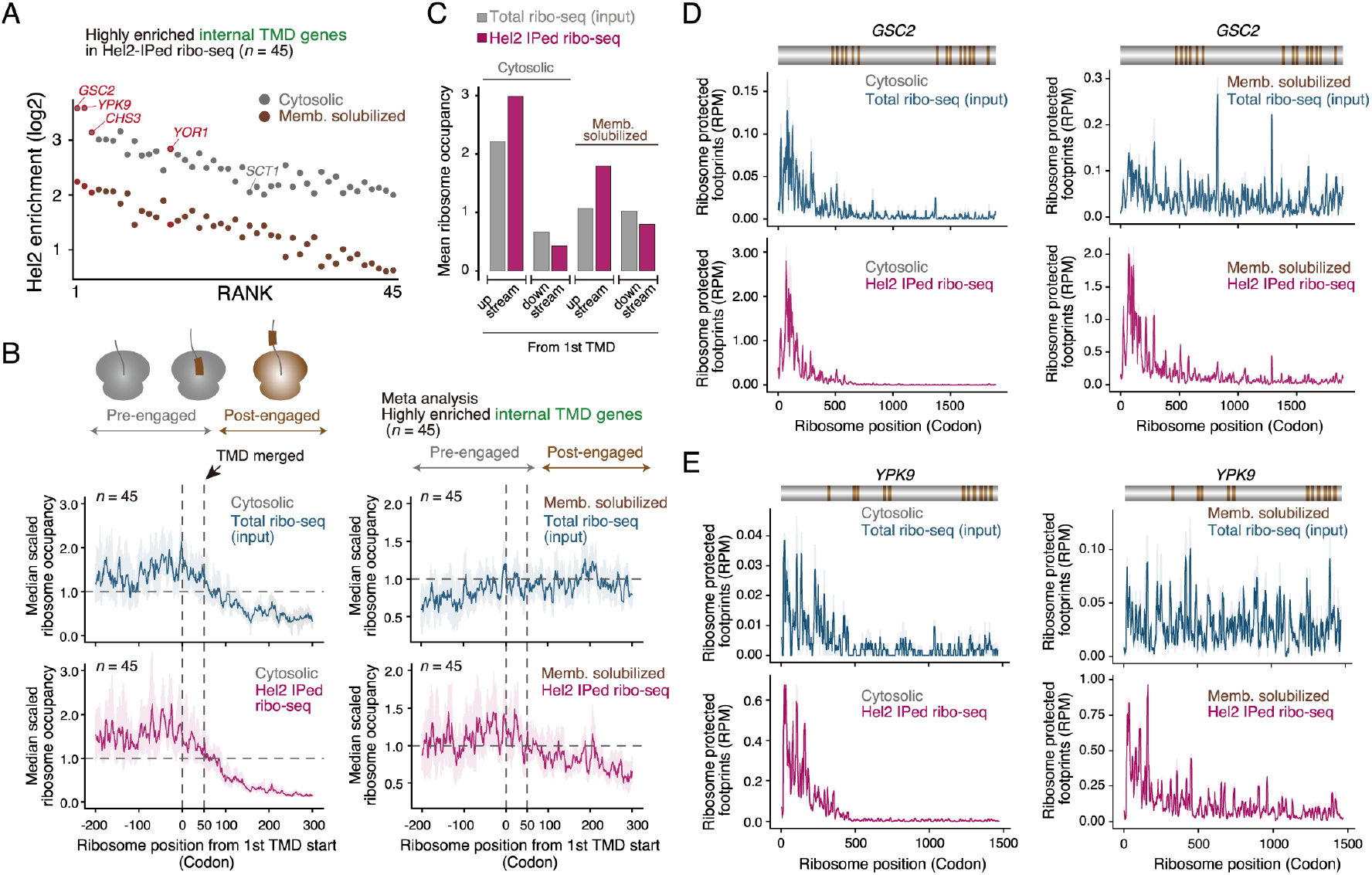
Hel2 monitors initial selection step for the secretory pathway. **(A)** Ranking of ER membrane-associated mRNAs encoding internal proteins. The top 45 ranked Hel2-enriched internal TMD genes were listed. Hel2 enrichment scores of individual transcripts from the cytosolic and membrane-solubilized fraction were plotted as gray and brown circles, respectively. The three top-ranked transcripts, *GSC2, YPK9*, and *CHS3*, are indicated in red, as is *YOR1*, a suggested RQC substrate (Lakshminarayan et al., 2020). The calculated values of each highly enriched internal TMD gene are listed in Supplemental Table 4. **(B)** Metagene plot of Hel2 enrichment score. Median-scaled ribosome occupancy of highly enriched internal TMD genes was plotted relative to the first TMD coding region of total ribo-seq and Hel2-IPed ribo-seq for the cytosolic and membrane-solubilized fractions (Chartron et al., 2016). **(C)** Ribosome occupancy up- and downstream of the first TMD coding region. Mean ribosome occupancy at each codon up- or downstream of the first TMD coding region was plotted for total ribo-seq and Hel2-IPed ribo-seq from cytosolic and membrane-solubilized fractions. **(D, E)** Ribosome density in the top-ranked ER membrane-associated mRNAs encoding internal proteins. RPMs of total ribo-seq and Hel2-IPed ribo-seq for (**D**) *GSC2* and (**E**) *YPK2* in the cytosolic and membrane-solubilized fractions. The membrane topology is indicated as above, in brown.

The Hel2-associated ribosomes on *GSC2*, the most enriched secretory mRNA encoding an internal protein in Hel2-IPed ribo-seq (**Figure 4A**), were highly concentrated in the 5’ region, within ~200 codons downstream of the initiation site (**Figure 4D**), whereas the first TMD was located at codon 464. Similar Hel2-recruitment was observed in *YPK9* and *CHS3* mRNAs, the second- and third-ranked Hel2-enriched mRNAs, respectively (**Figure 4E and S5A**). In addition, our top-ranked Hel2-enriched internal proteins included *YOR1* (**Figure 4A**), which may be a RQC-target (Lakshminarayan et al., 2020). On *YOR1* mRNA, Hel2 was again recruited during the early phase of elongation before membrane engagement (**Figure S5B**). Collectively, these observations indicated that Hel2 is predominantly recruited during the early phase of ribosome elongation rather than at engagement with the ER membrane.

### Hel2 targets secretory RNCs lacking SRP recognition

Although little is known about how SRP recognizes secretory RNCs before the targeting signal emerges from ribosome tunnel, SRP immediately binds to the secretory RNCs after initiation of translation in the SRP-preloading model (Chartron et al., 2016). In a representative SRP pre-recruited mRNA *VBA4*, Hel2 was recruited in the early phase of translation, similar to SRP-preloading system (**Figure S5C**). This finding suggested that Hel2 may monitor the initial selection step by SRP and capture problematic substrates. This hypothesis was tested by selective ribosome profiling of Hel2 under SRP-deficient conditions. The level of expression of *SRP72* was partially suppressed by replacement of its promoter, reducing the capacity of the SRP-mediated targeting system. Under the control of weak promoters (*AGX1p, BIT2p*, and *GIT1p*), Srp72 expression was decreased and a significant growth defect was observed (**Figure 5A**). Selective ribosome profiling was therefore performed using the *GIT1* promoter to suppress the expression of *SRP72* (**Figure S6A-D**). To assess the effects of SRP deficiency on Hel2-recruitment into secretory RNCs, we compared Hel2-enrichment among several cellular compartments. Notably, Hel2 enrichment in ER-targeted mRNAs, including SS and TMD coding sequences, was clearly increased under SRP-deficient conditions, whereas other components were not affected (**Figure 5B-C**). To further investigate the relationship between SRP and Hel2, we compared the Hel2-enrichment score with the SRP-dependent ER retention score of secretory mRNAs from a proximity-specific ribosome profiling dataset, performed using the Sec63-BirA system in the presence or absence of SRP, which is controlled by an auxin-inducible degradation system (Costa et al., 2018). Using these datasets, we estimated the effect of auxin treatment on ER retention, finding that the ER retention score of most Hel2-substrates was reduced after SRP depletion (**Figure 5D**). Collectively, these results strongly support our hypothesis that Hel2 targets secretory RNCs lacking SRP recognition.

**Figure 5.**
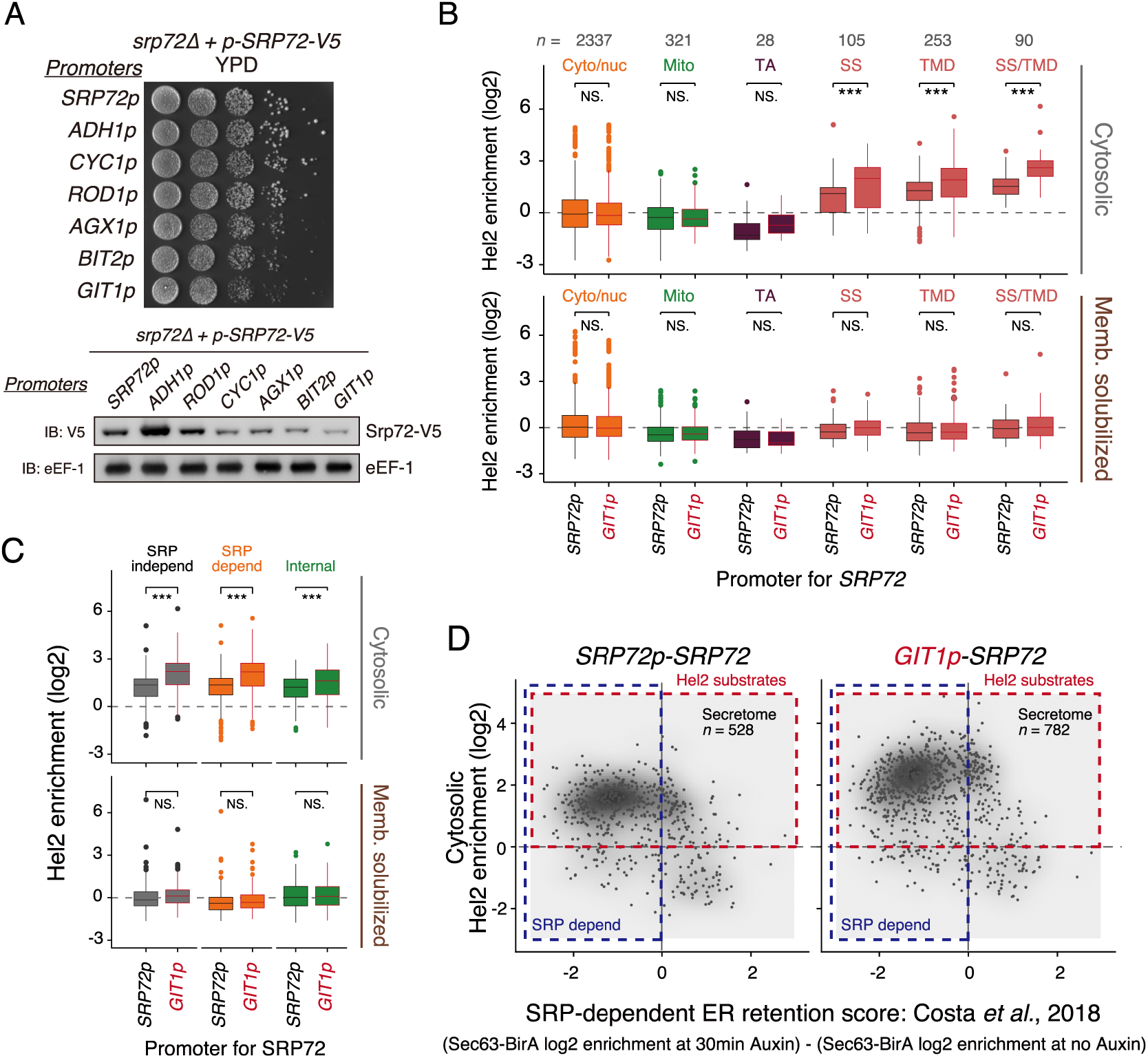
Hel2 targets to secretory RNCs lacking SRP recognition. **(A)** Upper panel: Growth defects due to suppression of *SRP72* by replacement with the indicated promoters. Lower panel: Immuno-blotting of Srp72-V5 with antibodies indicated as IB:X. **(B)** Box plot of Hel2 enrichment score of overall transcripts among cellular compartments (Chartron et al., 2016) for normal cells (*SRP72* promoter) and *SRP72*-deficient cells (*GIT1* promoter). Hel2 enrichment scores of individual transcripts were calculated by R package DEseq2. All Hel2-enrichment scores and P_adj_ for individual transcripts are shown in Supplemental Table 5. Significance was calculated by Student’s t-tests. NS, not significant, *P <0.05, ***P <0.001. **(C)** Box plot of Hel2 enrichment score of the transcripts encoding secretory proteins in normal cells (*SRP72* promoter) and *SRP72*-deficient cells (*GIT1* promoter). Hel2 enrichment scores of individual transcripts were calculated by R package DEseq2. The SRP dependence of individual transcripts was classified in (Costa et al., 2018). Significance was calculated by Student’s t-tests. NS, not significant, *P <0.05, ***P <0.001. **(D)** Scatter plot of cytosolic Hel2 enrichment versus SRP-dependent ER retention score. Hel2 enrichment scores of individual transcripts in normal cells (*SRP72* promoter) and *SRP72*-deficient cells (*GIT1* promoter) were calculated by R package DEseq2. SRP-dependent ER retention scores were calculated using previously published proximity-specific ribosome profiling datasets using the Sec63-BirA system in the presence or absence of SRP, which is controlled by an auxin-inducible degradation system (Costa et al., 2018). SRP-dependent ER retention scores were calculated by dividing Sec63-BirA enrichment (log2) with auxin treatment by Sec63-BirA enrichment (log2) without auxin treatment. GSM2836139, GSM2836140, GSM2836143, and GSM2836144 were used for the analyses.

### Hel2 prevents the mistargeting of secretory proteins into mitochondria

Although the mechanism to generate ribosome collisions during the early phase of translation elongation of secretory mRNAs is currently ill-defined, our observations suggest a preventive quality control system for secretory RNCs not recognized by SRP. Loss of SRP was found to lead to mitochondrial dysfunction caused by the mistargeting of secretory RNCs to mitochondria (Costa et al., 2018). Thus, we assessed whether Hel2 could cope with secretory RNCs lacking SRP recognition, which may be mistargeted to the mitochondria. Proximity mito-specific ribosome profiling (Tom20-BirA system) in the absence of SRP identified a list of secretory mRNAs mistargeted to the mitochondria (Costa et al., 2018). We calculated the Hel2-enrichment scores of 28 of these mistargeted mRNAs, which are enriched by more than 1 log2 on the mitochondrial outer membrane in the absence of SRP (**Figure S6E**: (Costa et al., 2018)). We found that these 28 mistargeted mRNAs were enriched on Hel2-associated ribosomes in the cytosolic fraction, with this enrichment clearly increased under SRP-deficient conditions (**Figure 6A**). To assess whether Hel2 prevents the mistargeting of secretory RNCs lacking SRP recognition, we utilized fluorescence microscopy to monitor the location of the ER-localized protein Sct1, which has been reported to be mistargeted to the mitochondria in the absence of SRP (Costa et al., 2018). Moreover, because *SCT1* mRNA was found to be highly enriched in Hel2-associated ribosome (**Figure 4A**), we employed Sct1-eGFP to evaluate mistargeting to the mitochondria. Although the *AGX1* promoter partially reduced the expression of *SRP72*, it did not affect the localization of Sct1-eGFP (**Figure 6B**), whereas additional *hel2* deletion resulted in Sct1-eGFP localizing to the mitochondria (**Figure 6B-C**). Moreover, although the combination of low expression of *SRP72* (*GIT1p-SRP72*) and deletion of *hel2* slightly reduced growth on glucose (fermentation) medium (**Figure 6D**), it markedly reduced growth on glycerol (respiratory) medium (**Figure 6D**), indicating that the mitochondrial defects induced by insufficient SRP are enhanced by the absence of Hel2.

**Figure 6.**
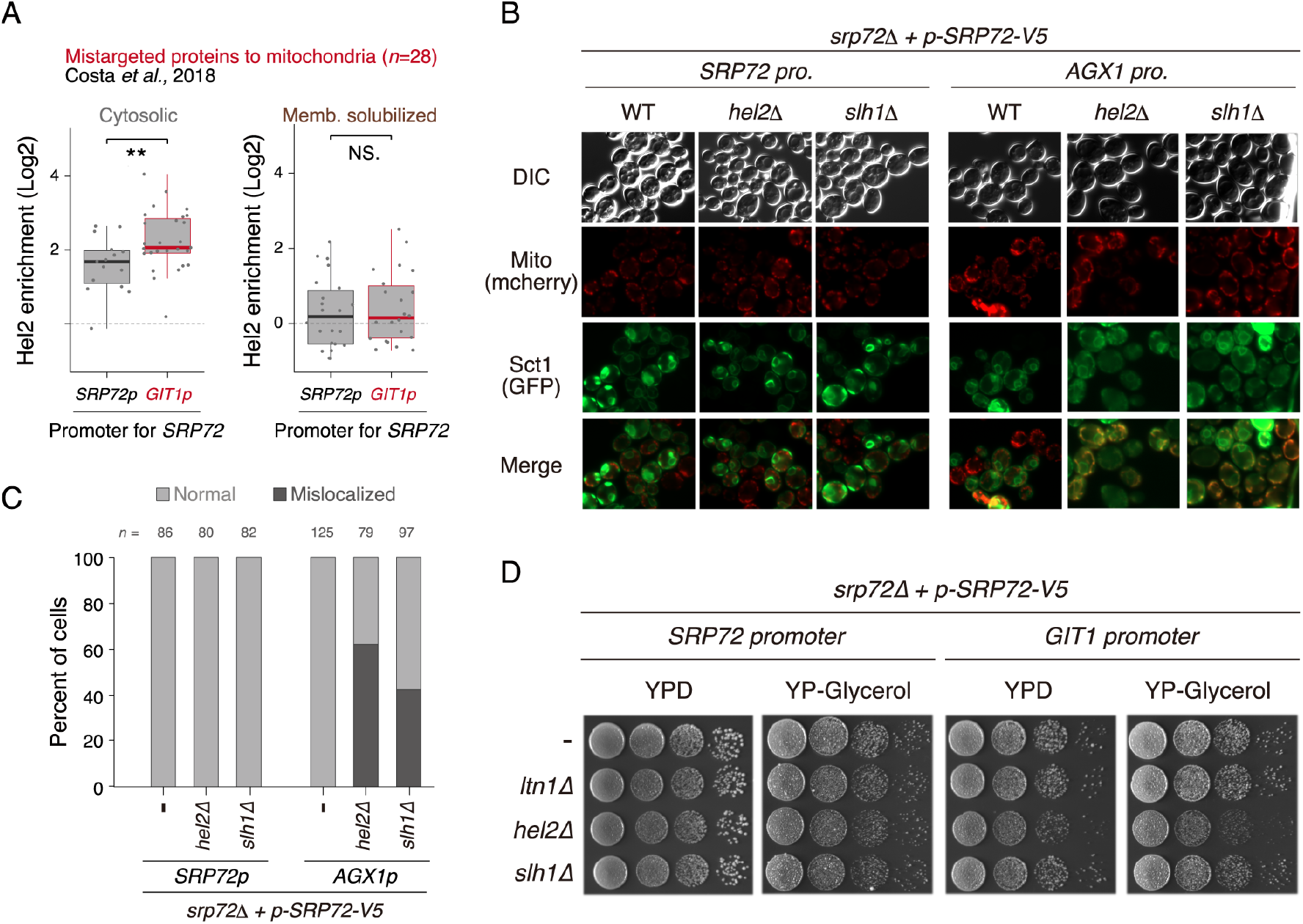
Hel2 prevents the mistargeting of secretory proteins into mitochondria. **(A)** Box plot of Hel2 enrichment scores of the proteins mistargeted to mitochondria in normal cells (*SRP72* promoter) and *SRP72*-deficient cells (*GIT1* promoter). Hel2 enrichment scores of individual transcripts were calculated by R package DEseq2. The proteins mistargeted to mitochondria were defined in **Figure S6E**. Significance was calculated by Student’s t-tests. NS, not significance; **P <0.01. **(B)** Subcellular distribution of Sct1-eGFP in various combinations of *SRP72* expression (*SRP72* or *AGX1* promoter) and *hel2* and *slh1* deletion analyzed by fluorescence microscopy. DIC: differential interference contrast; Mito: Mito-mCherry. **(C)** Quantification of the mistargeting rate in (B). Cells with Sct1-eGFP localized to the mitochondria were counted as incorrectly localized cells in various combinations of *SRP72* expression (*SRP72* or *GIT2* promoter) and *ltn1, hel2*, or *slh1* deletion. The ratios of normal to incorrectly localized cells were plotted. **(D)** Growth defects in various combinations of cells with *SRP72* expression (*SRP72* or *GIT2* promoter) and *ltn1, hel2*, or *slh1* deletion.

Because Hel2 is involved in ribosome-associated quality control (RQC), we further analyzed the effects of deletion of another RQT factor, *slh1*, on Sct1-eGFP localization (Matsuo et al., 2017; Matsuo et al., 2020). We found that the combination of *slh1* deletion and *SRP72* deficiency resulted in similar mislocalization of Sct1-eGFP (**Figure 6B-C**). However, this combination did not result in a growth defect (**Figure 6D**). Hel2 recognizes colliding ribosomes and induces three pathways, No-go mRNA decay (NGD), Smy2-Syh1-mediated mRNA decay, and the RQC pathway (D’Orazio et al., 2019; Hickey et al., 2020; Ikeuchi et al., 2019; Simms et al., 2017). Furthermore, the human Hel2 homolog ZNF598 was found to inhibit translation initiation via GIGYF2-4EHP system on colliding ribosomes in mammalian cells (Hickey et al., 2020). By contrast, Slh1 directly splits the colliding ribosomes to induce the RQC pathway after the ubiquitination of uS10 by Hel2 (Matsuo et al., 2020), but is not involved in the NGD or Smy2-Syh1-mediated mRNA decay pathways. These findings indicate that mislocalized-secretory RNCs could be suppressed by several pathways induced by the Hel2-mediated quality control system, including the RQC pathway.

## Discussion

Taken together, our findings suggest a model by which Hel2 plays a crucial role in the triage system to prevent the mistargeting of secretory proteins. The transcripts encoding secretory proteins are targeted to the ER by the SRP-mediated system (Guna and Hegde, 2018; Zhang and Shan, 2014). Although this system is very accurate, a small population of transcripts may be unrecognized by SRP, resulting in their aggregation in the cytosol or their being mistargeted to mitochondria (Costa et al., 2018). Selective ribosome profiling revealed that the mRNAs encoding secretory proteins are particularly enriched in Hel2-IPed ribo-seq, suggesting that secretory RNCs lacking SRP recognition could induce translational arrest by as-yet undefined mechanisms, generating a ribosome traffic jam recognized by Hel2 as a problematic event in translation. The force-quit of translation by Hel2 during the early phase of elongation may be optimal for coping with potentially mistargeted membrane proteins because hydrophobic regions have not yet been translated. This preventive action may greatly contribute to avoiding the production of harmful nascent peptides containing hydrophobic regions. Indeed, the products derived from this preventive mechanism seemed to be less toxic because the lack of Ltn1 had no effect on the growth of SRP-insufficient cells. RQT factors including Slh1 are targeted only to the leading ribosome at the collision site, inducing its disassembly (Juszkiewicz et al., 2020; Matsuo et al., 2020). Because the trailing ribosomes can still elongate after the leading ribosome is dissociated by the RQT complex, this dissociation may provide the trailing ribosomes with a second opportunity to engage with the ER. Collectively, these findings provide novel insights into the risk management of membrane protein biogenesis for maintaining cellular protein homeostasis.

Our selective ribosome profiling showed that Hel2 apparently binds to all types of secretory RNCs, including SRP-dependent and −independent proteins. SRP is the main factor responsible for ER targeting system of secretory RNCs, except for tail-anchored proteins, but other routes to the ER membrane have been suggested. Although genetic dependence on SRP is a good indicator of its physical engagement with the ER membrane, SRP independence cannot be equated with the lack of SRP engagement under normal conditions. Indeed, SRP-selective ribosome profiling showed that SRP binds to nearly all secretory RNCs that are co-translationally targeted to the membrane, including SRP-dependent and SRP-independent proteins (Chartron et al., 2016), similar to the results of Hel2-selective ribosome profiling. Furthermore, the SND pathway was recently identified as an alternative route for ER targeting, independent of the SRP-mediated targeting system (Aviram et al., 2016). The SND pathway specifically recognizes substrates with central TMDs and partially suppresses the loss of SRP (Aviram et al., 2016), suggesting that it may act as a backup system for SRP-mediated targeting. This finding explains why SRP independence cannot be equated with lack of SRP engagement under normal conditions, in that secretory RNCs that failed to engage with the ER via SRP could be recaptured by the alternative SND pathway, resulting in correct engagement with the ER even in the absence of SRP. Although Hel2 recognizes secretory RNCs lacking SRP recognition as well as the SND pathway, its recognition mechanism seems to differ from the SND pathway because Hel2 is recruited only during the early phase of translation elongation. However, its recognition mechanism has yet to be determined. Future studies should focus on the selectivity of substrates by Hel2 and further analyze the relationship between the SND-mediated targeting system and the Hel2-mediated quality control system, which may provide better understanding of the fate of secretory proteins lacking SRP recognition.

The present study found that Hel2 recognized unengaged secretory RNCs due to their lack of SRP recognition, and contributed to preventing the mistargeting of secretory proteins into the mitochondria. However, the mechanistic details of Hel2-mediated quality control are not yet fully understood. During the initial step of the RQC pathway, Hel2 recognizes the colliding ribosomes and ubiquitinates ribosomal protein uS10, triggering ribosome dissociation into its subunits (Matsuo et al., 2017). This non-canonical ribosome dissociation results from an energy-consuming reaction by the RQT complex, which consists of RNA helicase Slh1, ubiquitin-binding protein Cue3, and zinc finger protein Rqt4 (Matsuo et al., 2020). Although the deletion of *slh1* under SRP-deficient conditions led to the mistargeting of Sct1 into the mitochondria, a significant growth defect was not observed under respiratory conditions, indicating that the RQC pathway partially contributes to suppressing the mistargeted secretory RNCs. Hel2 is a master regulator, inducing several pathways on the colliding ribosomes, whereas Slh1 is involved only in the RQC pathway. The two additional Hel2-mediated mRNA degradation pathways have been identified, Cue2-mediated No-Go-decay and Smy2-Syh1-mediated mRNA decay (D’Orazio et al., 2019; Hickey et al., 2020; Ikeuchi et al., 2019; Simms et al., 2017). Smy2 and Syh1 are yeast homologs of GIGYF2. Furthermore, the human Hel2 homolog ZNF598 was found to inhibit translation initiation via the GIGYF2-4EHP system on the colliding ribosomes in mammalian cells (Hickey et al., 2020). Although 4EHP is not conserved in yeast, Syh1 interacts with Eap1, an inhibitor of eIF4E function (Sezen et al., 2009), suggesting that a similar mechanism in yeast might suppress ribosome collision and protein production. In contrast to *slh1* deletion, *hel2* deletion in combination with *SRP72*-deficiency resulted in a severe growth defect under respiratory conditions, suggesting that Hel2 induces additional pathways, including NGD, Smy2-Syh1 mediated mRNA decay, and initiation control, to cope with mistargeting of secretory RNCs. Further analysis of these downstream pathways will uncover the mechanistic details of Hel2-mediated quality control in secretory pathways.

The lack of SRP on the secretory RNCs allows for Hel2 recruitment, however, the mechanism underlying this process remains largely unknown. The many secretory RNCs were recognized by SRP before the targeting signal emerges from ribosome tunnel (Chartron et al., 2016). But why and how SRP recognizes secretory RNCs in such an early phase of translation remains enigmatic. These questions should be addressed in the future studies to understand the recognition mechanism of secretory RNCs by SRP and Hel2.

## Author Contributions

Experiments were designed and data were interpreted by Y.M. and T.I.; Y.M. performed the experiments and analyzed ribosome-profiling datasets; the manuscript was written by Y.M. and T.I.

## Acknowledgments

Part of this work was performed at the Vincent J. Coates Genomics Sequencing Laboratory at UC Berkeley, supported by NIH S10 Instrumentation Grant OD018174. Computations were partially performed on the NIG supercomputer at the ROIS National Institute of Genetics. This study was supported by a Grant-in-Aid for Scientific Research (KAKENHI) from the Japan Society for the Promotion of Science (grant numbers 18H03977 to T.I. and 19K06481 to Y.M.) and by Research Grants in Mochida Memorial Foundation for Medical and Pharmaceutical Research (to Y.M.).

## SATAR Methods

### RESOURCE AVAILABILITY

#### Lead Contact

Please direct any requests for further information or reagents to the lead contact, Toshifumi Inada.

#### Materials Availability

Plasmids and genetically engineered yeast strains that have been generated as part of this study are available upon request.

#### Data and Code Availability

The sequencing data for ribosome profiling experiments have been deposited in NCBI’s Gene Expression Omnibus and are accessible through GEO series accession number GSE156535. The scripts written by R for data analysis are available upon request. Original images used for the figures have been deposited to Mendeley Data: http://dx/doi.org/10.17632/2ngd5vgw7f.1

### EXPERIMENTAL MODEL AND SUBJECT DETAILS

Derivatives of the *S. cerevisiae* W303 strain were used for the experiments presented. The *S. cerevisiae* strains used in this study are listed in the **Key Resources Table**. Gene disruption and C-terminal tagging were performed as described (Janke et al., 2004; Longtine et al., 1998). The knock-in or −out of transformants were confirmed by PCR, and then again confirmed by western blotting or RT-PCT, respectively. Most of the yeast strains were grown in Yeast Extract-Peptone-Dextrose (YPD) medium at 30 °C. The details of the culture conditions for individual experiments were described in the METHOD DETAILS section.

### METHOD DETAILS

#### Plasmid constructs

All recombinant DNA techniques were performed according to standard procedures using *E. coli* DH5α for cloning and plasmid propagation. All cloned DNA fragments generated by PCR amplification were verified by sequencing. Plasmids used in this study are listed in the **Key Resources Table.**

#### Selective ribosome profiling

##### Preparation of cytosolic and membrane-solubilized fractions

The genomically tagged yeast strain *HEL2-FTP* (Flag-TEV-ProteinA) was cultured in 6 L YPD medium until mid-log phase (OD600 = 0.5 ~ 0.6). The cells were harvested by centrifugation at 6,000 rpm for 2 minutes at 30 °C using a JLA-8.1000 rotor (Beckman Coulter). Each harvested cell pellet was immediately frozen in liquid nitrogen, and then ground in liquid nitrogen using a mortar and pestle. The powder was resuspended in lysis buffer (50 mM Tris, pH 7.5, 100 mM NaCl, 10 mM MgCl2, 1 mM DTT, with or without 1 % (vol/vol) Triton X-100) containing 100 μg/ml cycloheximide and 100 μg/ml tigecycline. Whole cell lysates were centrifuged at 40,000 g for 30 min at 4°C, and the supernatant fractions was used for the following steps.

##### Preparation of ribosome fraction for total ribo-seq (input)

The supernatant fraction containing 10 μg of total RNA was treated with 12.5 units of RNase I (Epicentre) at 23 °C for 45 min (strong RNase I treatment), followed by sedimentation through a 1 M sucrose cushion. Although mild RNase I treatment were performed in the preparation for Hel2-IPed ribo-seq to separate individual ribosomes, it was skipped in the total ribo-seq preparation. The strong RNase I treatment is enough to generate the individual ribosome footprints. The ribosome-protected mRNA fragments were extracted with TRIzol reagent (Life Technologies) and used for library preparation.

##### Preparation of ribosome fraction for Hel2-IPed ribo-seq

To obtain the cytosolic fraction, 1% (vol/vol) Triton X-100 was added to the supernatant fraction after initial centrifugation. The supernatant fraction containing 80 mg of total RNA was partially digested with 250 units of RNase I (Epicentre) at 4 °C for 1 hour (mild RNase I treatment), followed by incubation with IgG conjugated Dynabeads (Invitrogen) for 1 hour at 4 °C for binding of Hel2 to these beads. The IgG beads were washed with lysis buffer, followed by TEV protease cleavage. The eluate containing 10 μg of total RNA was again digested with 12.5 units of RNase I (Epicentre) at 23 °C for 45 min (strong RNase I treatment), and the reaction was stopped by adding TRIzol reagent (Thermo Fisher Scientific), followed by extraction of the ribosome-protected mRNA fragments.

##### Library preparation

The extracted RNA was size-selected from 15% denaturing PAGE gels, cutting between 26–34 nt. Libraries were prepared as described, with several modifications (Ingolia et al., 2012). The linker DNA consisted of 5’-(Phos)NNNNNIIIIITGATCGGAAGAGCACACGTCTGAA(ddC)-3’ where (Phos) indicates 5’ phosphorylation and (ddC) indicates a terminal 2’, 3’-dideoxycytidine. The Ns and Is indicate a unique molecular identifier (UMI) to eliminate PCR duplication and multiplexing, respectively. The linkers were pre-adenylated with a 5’ DNA adenylation kit (NEB) and used for the ligation reaction. An oligonucleotide, 5’-(Phos)NNAGATCGGAAGAGCGTCGTGTAGGGAAAGAG(iSp18)GTGACTGGAGTTC AGACGTGTGCTC-3’, where (Phos) indicates a 5’ phosphorylation site and Ns indicate a random sequence, was used for reverse transcription. PCR was performed using the oligonucleotides 5’-AATGATACGGCGACCACCGAGATCTACACTCTTTCCCTACACGACGCTC-3’ and 5’-CAAGCAGAAGACGGCATACGAGATJJJJJJGTGACTGGAGTTCAGACGTGTG-3’, where Js indicate the reverse complement of the index sequence discovered during Illumina sequencing. The libraries were sequenced on a HiSeq 4000 (Illumina).

#### Analysis of ribosome profiling

Sequencing reads were de-multiplexed and stripped of 3’ linker sequences using FASTX-toolkit v0.0.14. The UMI, which can serve to remove PCR duplications generated during library preparation, were extracted by UMI-tools v1.0.1 (Smith et al., 2017). The reads were first filtered by mapping to Bowtie Index, composed of non-coding RNA genes, using Bowtie2 v2.2.5 (Langmead and Salzberg, 2012). Reads were mapped to the genome using Tophat v2.1.1 (Kim et al., 2013). Only uniquely-mapping reads from the final genomic alignment were used for subsequent analysis. The position of the A site from the 5’-end of the reads at the initiation codon was estimated based on the length of each footprint using the plastid v0.4.7 (Dunn and Weissman, 2016). The mapped read counts were calculated by plastid v0.4.7. using two modes: 5’-end assignment for A site positioning and center weighting for ribosome positioning. The 28–31 nt long reads were regarded as ribosome-protected mRNA fragments, and the offsets used for 5’-end assignment were 15 for 28 nt and 16 for 29–31 nt. Transcript enrichment was computed by R package “DESeq2” (Love et al., 2014). Transcripts with fewer than 25 reads were omitted from transcript enrichment analysis. Individual ORFs were classified as described (Ast et al., 2013; Chartron et al., 2016; Costa et al., 2018). The ribosome occupancy was calculated by R v3.3.2 as the ratio of reads at a given codon or nucleotide position to the average reads per codon on individual transcripts. The Hel2-enrichment score at codon resolution was calculated by dividing the RPM of Hel2-IPed ribo-seq by the RPM of total ribo-seq (input) using R v3.3.2. For metagene analysis, mean or median-scaled ribosome occupancy and median-scaled Hel2-enrichment at given codons were calculated by R v3.3.2. First TMD start sites were as in (Chartron et al., 2016). Transcripts with <0.5 average reads per codon were omitted from metagene analysis. Gene ontology enrichment analysis in the cellular component was performed as described (http://geneontology.org). In the selection of highly enriched internal TMD genes in Hel2-IPed ribo-seq, transcripts with fewer than 100 reads were omitted. The analysis of proximity-specific ribosome profiling used previously published GSM2836139, GSM2836140, GSM2836143, GSM2836144, GSM2836161, GSM2836162, GSM2836163, and GSM2836164 datasets (Costa et al., 2018).

#### In vitro translation of SDD1 mRNA

*SDD1 reporter* mRNA was produced using the mMessage mMachine Kit (Thermo Fischer) and used in a yeast cell-free translation extract from *uS10-3HA ski2D* cells. This yeast translation extract was prepared and *in vitro* translation was performed essentially as described before ^49^. The cells were grown in YPD medium to an OD600 of 1.5-2.0, washed with water, 1% KCl and finally incubated with 10 mM DTT in 100 mM Tris, pH 8.0 for 15 min at room temperature. To generate spheroplasts, 2.08 mg zymolyase per 1 g of cell pellet was added in YPD 1 M sorbitol and incubated for 75 min at 30°C. Spheroplasts were then washed three times with YPD 1 M sorbitol and once with 1 M sorbitol and lysed as described before ^49^ with a douncer in lysis buffer (20 mM HEPES pH 7.5, 100 mM KOAc, 2 mM Mg(OAc)2, 10% Glycerol, 1 mM DTT, 0.5 mM PMSF and complete EDTA-free protease inhibitors (GE Healthcare)). From the lysate, a S100 fraction was obtained by low-speed centrifugation followed by ultracentrifugation of the supernatant. The S100 was passed through a PD10 column (GE Healthcare). *In vitro* translation was performed at 17 °C for 60 min using great excess of template mRNA (38 μg per 415 μl of extract) to prevent degradation of resulting stalled ribosomes by endogenous response factors. For Hel2-supplemented *in vitro* translation reaction, 32 nM Hel2 were supplied to the reaction.

#### Purification of RNCs on the SDD1 mRNA

The stalled RNCs on the *SDD1* mRNA were affinity purified using the His-tag on the nascent polypeptide chain and magnetic IMAC-beads. After *in vitro* translation at 17 °C for 60 min the extract was applied to Dynabeads^™^ (Invitrogen) for His-Tag isolation and pulldown for 15 min at 4 °C. The beads were washed with a lysis buffer 300 (50 mM HEPES pH 7.5, 300 mM KOAc, 10 mM Mg(OAc)2, 0.01% NP-40 and 5 mM β-Mercaptoethanol) and eluted in the elution buffer (50 mM HEPES pH 7.5, 100 mM KOAc, 10 mM Mg(OAc)2, 0.01% NP-40, 5 mM β-Mercaptoethanol) containing 300 mM imidazole.

#### Sucrose density gradient sedimentation analysis

Samples were layered onto 10–50% (w/v) sucrose gradients in 25 mM Tris, pH 7.5, 100 mM NaCl, and 10 mM MgCl2, followed by centrifugation at 40,000 rpm for 1 hour 45 minutes at 4 °C using an SW-41Ti rotor (Beckman Coulter). The absorbance of the gradient at 254 nm relative to the length of the gradient was monitored by gradient station (Biocomp).

#### Immunoblotting

Proteins were separated by 10% Nu-PAGE Bis-Tris gels, transferred to PVDF membranes (Millipore; IPVH00010), and blocked with 5% skim milk. The blots were incubated with anti-V5 (AbD Serotec: MCA1360) or anti-eEF1 (homemade) antibodies, followed by incubation with secondary antibodies conjugated to horseradish peroxidase (HRP). Bands were detected by ImageQuant LAS4000 (GE Healthcare).

#### Fluorescence microscopy

Exponential growing cells expressing Sct1-eGFP and Mito-mCherry were grown at 30°C for 1 hour in selective SC medium containing galactose. The cells were harvested by centrifugation, followed by fluorescence microscopy at room temperature using a microscope (Axio Imager 2 with EC Plan-NEOFLUAR 100x/1.30: ZEISS).

##### Quantification and statistical analysis

Meta-analyses were plotted mean- or median-scaled score as indicated in Figures (Figure 1H, Figure 2C-D, Figure 3D-E, Figure 4B, Figure S3I, and Figure S4E). The Spearman’s correlations (p) of scatter plots (Figure S2B, Figure S3G, and Figure S6A-D) were calculated by R. Significance of Box plots (Figure 5B-C, Figure 6A) were calculated by Student’s t-test.

## Supplementary Information

### Supplemental Figure Legends

**Figure S1.**
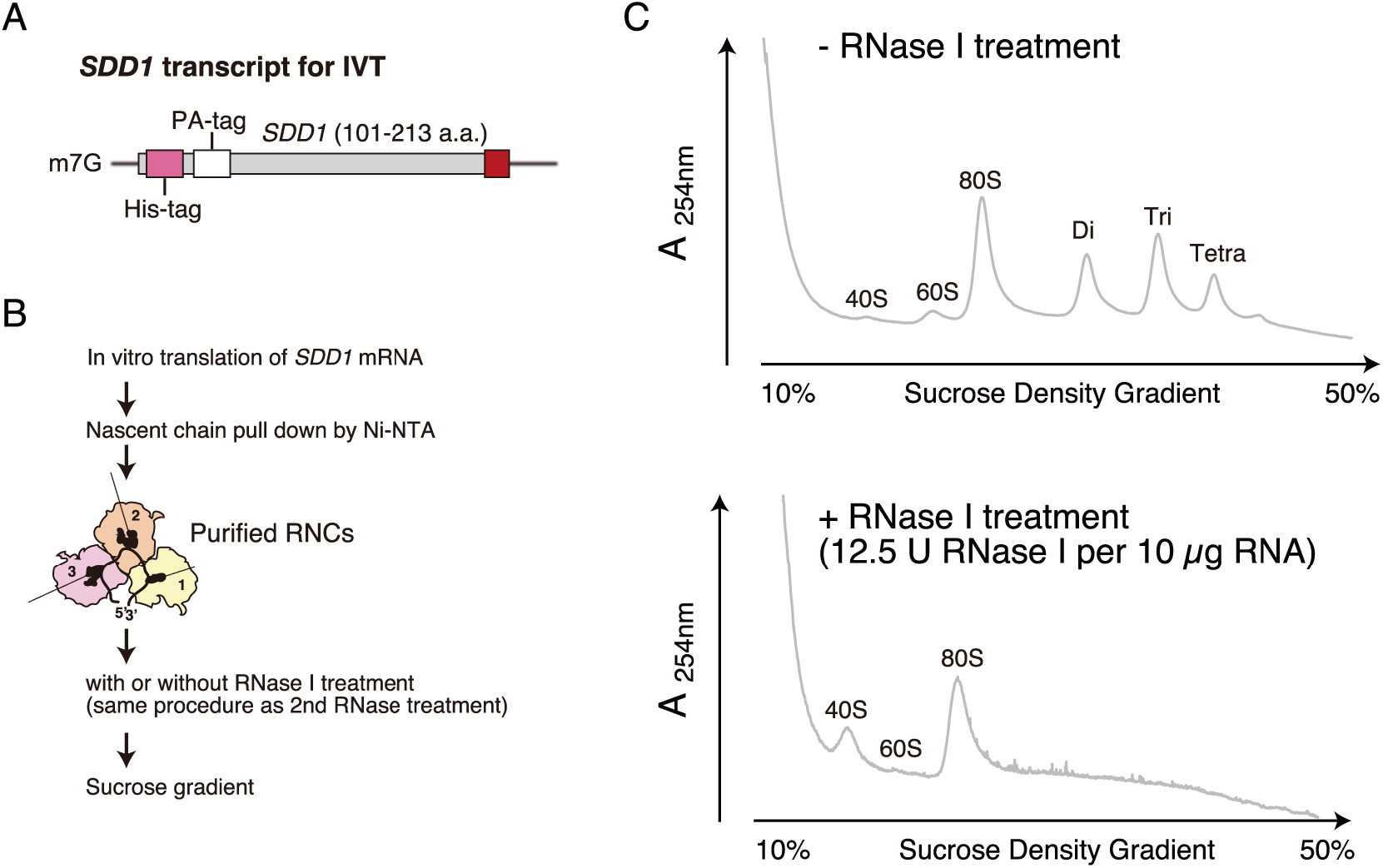
RNase resistant level of Hel2-associated colliding ribosomes on the *SDD1* mRNA: related to Figures 1. **(A)** Schematic drawing the *SDD1* model transcript. **(B)** Scheme of the *in vitro* reconstitution of the colliding ribosomes and their RNase I treatment. **(C)** RNCs were prepared by IVT as described in (B). After RNase I treatment, RNCs were separated by sucrose density gradient and detected ribosomes by UV absorbance at 254 nm wavelength.

**Figure S2.**
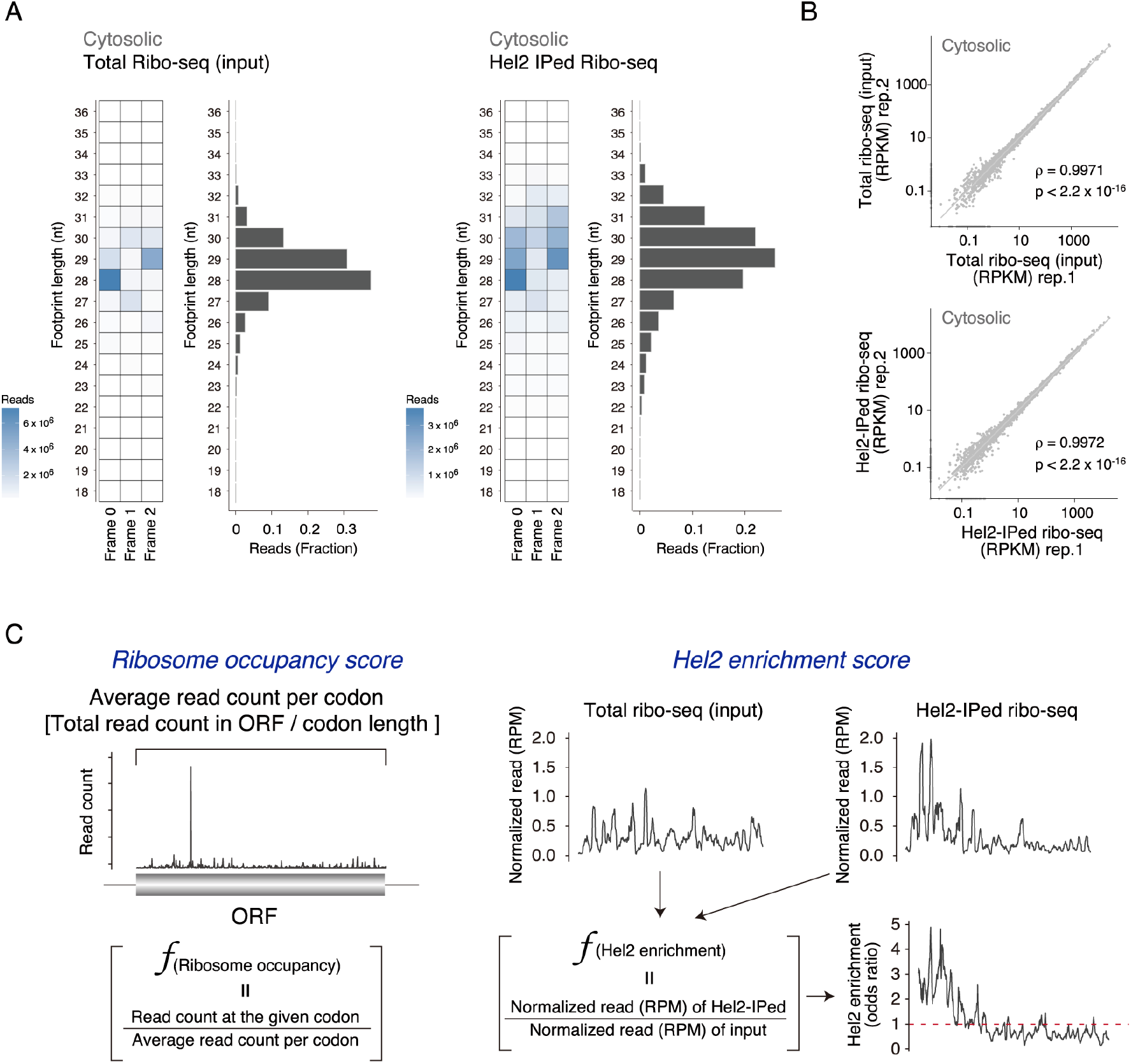
Selective ribosome profiling with the cytosolic fraction: related to Figures 1 and 2. **(A)** Codon periodicity and length distribution of ribosome-protected mRNA fragments. Heatmap of genome mapped reads at different lengths and frames with length distribution. Left panel: total ribo-seq. Right panel: Hel2-IPed ribo-seq. **(B)** Correlations of the sum of ribosome-protected mRNA fragments among biological replicates. Left panel: total ribo-seq. Right panel: Hel2-IPed ribo-seq. Spearman’s rank correlation ρ and P-value were calculated using R v3.3.2 software. **(C)** Calculations of ribosome occupancy and Hel2-enrichment scores.

**Figure S3.**
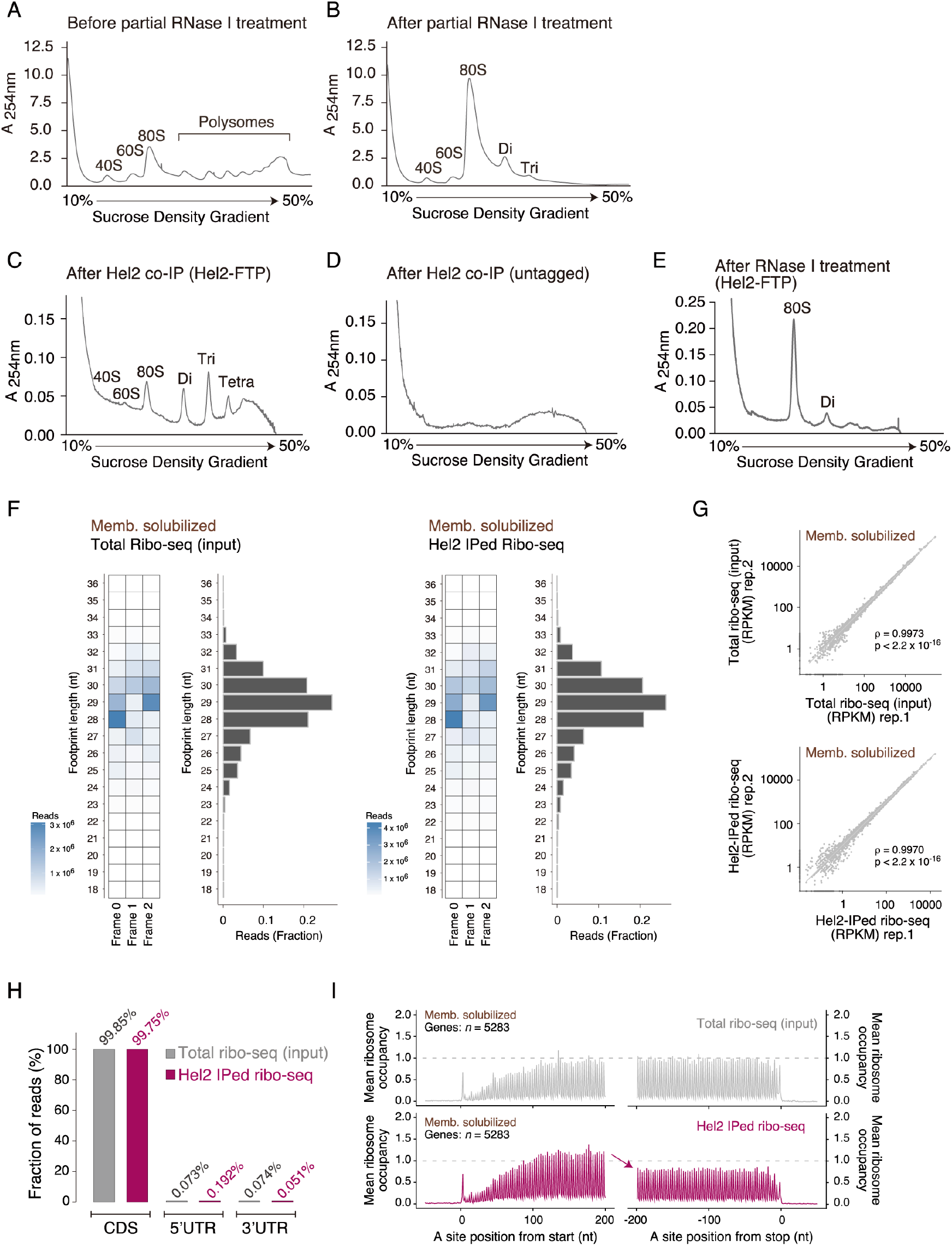
Selective ribosome profiling with membrane-solubilized fraction: related to Figure 3. **(A-E)** Sucrose density gradient sedimentation analysis. The disrupted cell powder was resuspended in lysis buffer and centrifuged at 40,000 x g for 30 minutes at 4 °C **(A)**. The resulting supernatant was partially digested with RNase I for 1 hour at 4 °C **(B)**, and Hel2-associated ribosomes were purified by immunoprecipitation **(C)**. The supernatant prepared from untagged-yeast strain was used as a negative control for immunoprecipitation **(D)**. The purified Hel2-associated ribosomes were digested with RNase I for 45 minutes at 23 °C to generate the ribosome-protected mRNA fragments **(E)**. Fractions obtained during each step of Hel2-associated ribosome purification were analyzed by 10–50% (w/v) sucrose density gradient centrifugation. **(F)** Codon periodicity and length distribution of ribosome-protected mRNA fragments. Heatmap of genome mapped reads at different lengths and frames with length distribution. Left panel: total ribo-seq. Right panel: Hel2-IPed ribo-seq. **(G)** Correlation of the sum of ribosome-protected mRNA fragments among biological replicates. Upper panel: total ribo-seq. Lower panel: Hel2-IPed ribo-seq. Spearman’s rank correlation p and P-values were calculated using R v3.3.2 software. **(H)** Fractions of reads mapping to each reading frame in the coding sequence (CDS), the 5’-untranslated region (UTR), and the 3’-UTR for total ribo-seq and Hel2-IPed ribo-seq. **(H)** Metagene plot of ribosome (nt) occupancy. Ribosome occupancy scores were calculated as the ratio of reads at each nucleotide position to average reads per codon on the individual ORF (**Figure S2C**). Average ribosome occupancy was plotted around the start and stop codons at A site for total ribo-seq and Hel2-IPed ribo-seq. The 28 nt length reads and the transcripts containing more than 0.5 reads per codon were included in the analyses. The offset was 15.

**Figure S4.**
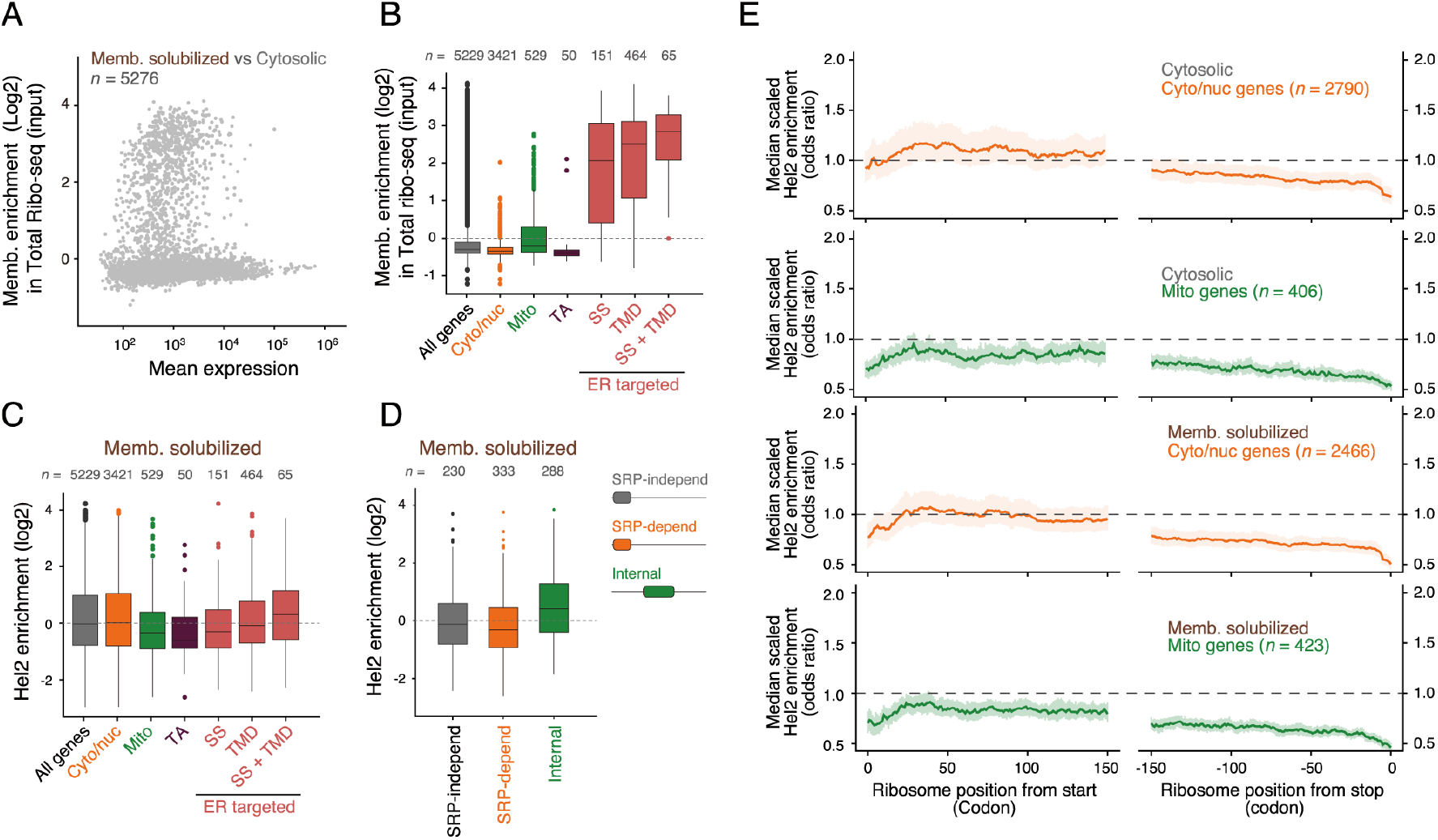
Hel2 does not bind to secretory RNCs that are engaged with ER: related to Figure 3. **(A)** Scatter plot of membrane enrichment, plotted as fold change, of total ribo-seq between membrane-solubilized and cytosolic fractions versus mean expression. The membrane enrichment score of individual transcripts was calculated by R package DEseq2. Analyses were restricted to transcripts mapped with more than 25 reads. All membrane-enrichment scores for individual transcripts are shown in Supplemental Table 2. **(B)** Box plot of membrane enrichment scores of overall transcripts among cellular compartments. Individual transcripts were classified into cellular compartments as described (Chartron et al., 2016). **(C)** Box plot of Hel2 enrichment scores of overall transcripts among cellular compartments. Hel2-enrichment scores for individual transcripts were calculated by R package DEseq2. All Hel2-enrichment scores of individual transcripts are shown in Supplemental Table 3. Individual transcripts were classified into cellular compartments as described (Chartron et al., 2016). **(D)** Box plot of Hel2 enrichment scores of transcripts encoding secretory proteins. The SRP dependence of individual transcripts was determined as described (Costa et al., 2018). Internal proteins have a first TMD after the initial 60 amino acids. **(E)** Metagene plot of Hel2 enrichment scores. Hel2 enrichment scores were calculated by dividing the RPM of Hel2-IPed ribo-seq by RPM of total ribo-seq (**Figure S2C**). Median-scaled Hel2 enrichment scores of cyto-nuclear genes and mitochondrial genes were plotted around the start and stop codons for cytosolic and membrane-solubilized fractions. The transcripts containing more than 0.5 reads per codon were analyzed. Individual transcripts were classified into cellular compartments as described (Chartron et al., 2016).

**Figure S5.**
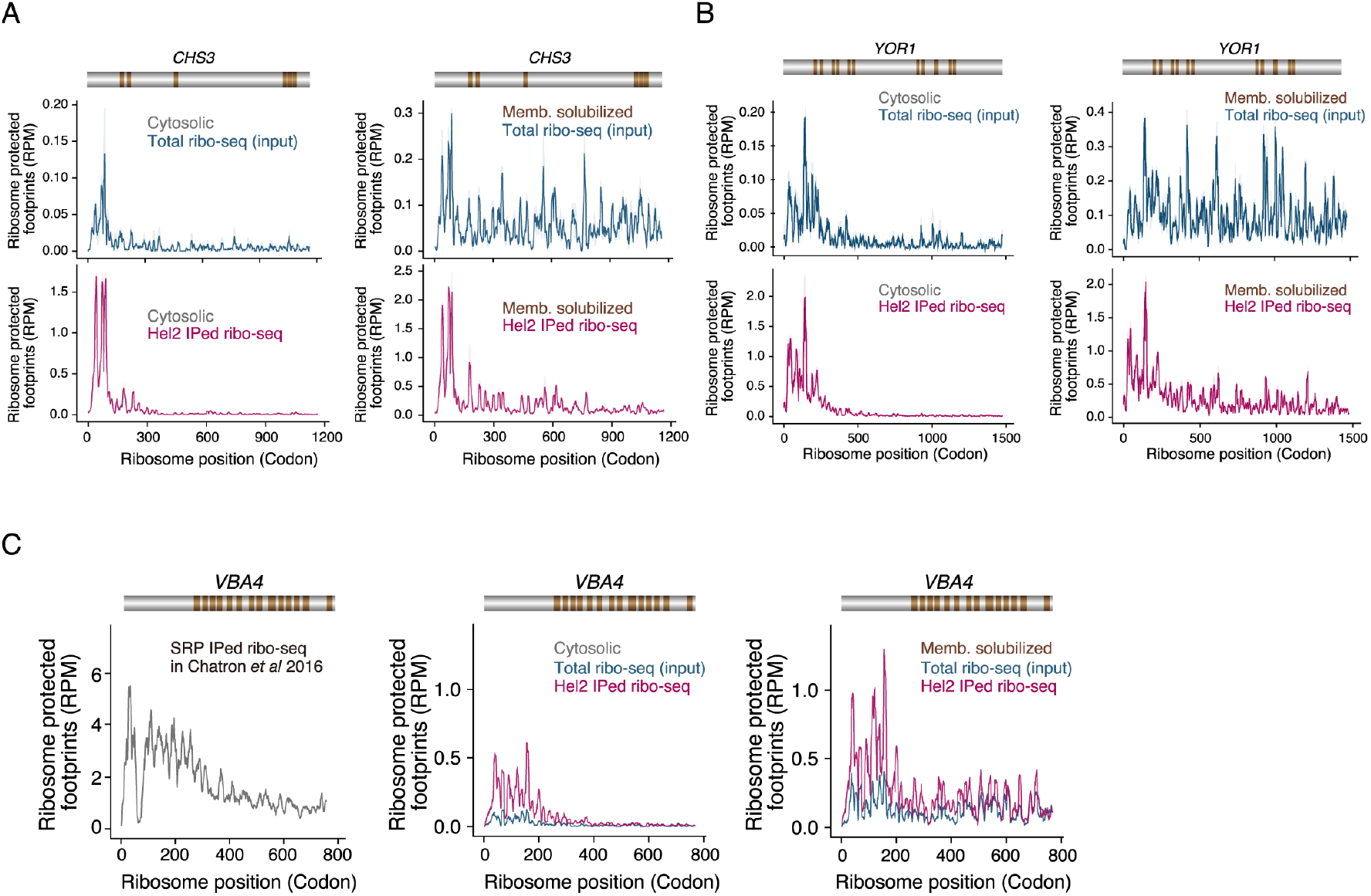
Ribosome density on individual genes: related to Figure 4. **(A, B, C)** The RPM of total ribo-seq and Hel2-IPed ribo-seq were plotted on *CHS3* (A), *YOR1* (B) for cytosolic and membrane-solubilized fraction. Membrane topology is shown as above, in brown. **(C)** The comparison of RPM between SRP- and Hel2-IPed ribo-seq on *VBA4* mRNA. The RPM of SRP IPed ribo-seq were calculated using previously published dataset (Chartron et al., 2016): GSM1919467.

**Figure S6.**
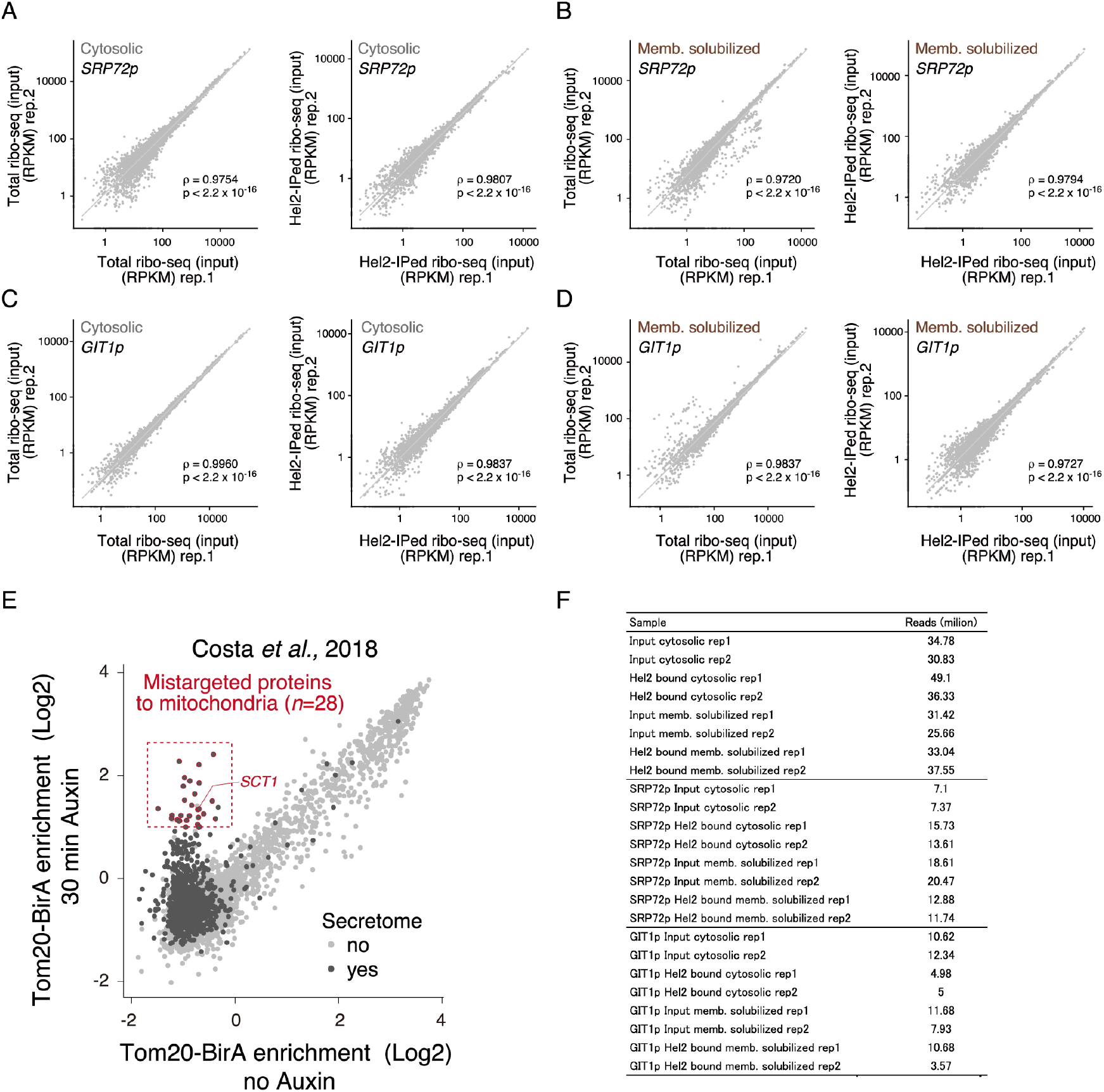
Selective ribosome profiling under the SRP-deficient condition: related to Figure 5 & 6. **(A-D)** Left panel: total ribo-seq. Right panel: Hel2-IPed ribo-seq. Spearman’s rank correlation: ρ, and P-values were calculated using R v3.3.2 software. **(E)** Scatter plots of Tom20-BirA enrichment (log2) in SRP-depleted (30 min Auxin) versus normal cells (no Auxin), calculated by R package DEseq2 using the previously published dataset (Costa et al., 2018): GSM2836161, GSM2836162, GSM2836163, and GSM2836164. **(F)** Coverage of datasets analyzed in this study. Indicated numbers of ribosome-protected mRNA fragments are mapped to coding sequences after the removal of PCR duplicates.

**Supplemental Table 1.** Hel2 enrichment in the cytosolic fraction: related to Figure 2.

**Supplemental Table 2.** Membrane enrichment: related to Figures 3 and S3.

**Supplemental Table 3.** Hel2 enrichment in membrane-solubilized fraction: related to Figures 3 and S3.

**Supplemental Table 4.** Highly enriched internal TMD genes: related to Figure 4.

**Supplemental Table 5.** Differences in Hel2 enrichment between normal and SRP72-deficient cells: related to Figure 5.

